# Harnessing the Intrinsic Photochemistry of Isoxazoles for the Development of Chemoproteomic Crosslinking Methods

**DOI:** 10.1101/2021.12.23.473999

**Authors:** Marshall G. Lougee, Vinayak Vishnu Pagar, Hee Jong Kim, Samantha X. Pancoe, Robert H. Mach, Benjamin A. Garcia, E. James Petersson

## Abstract

Photo-crosslinking is a powerful technique for identifying both coarse- and fine-grained information on protein binding by small molecules. However, the scope of useful functional groups remains limited, with most studies focusing on diazirine, aryl azide, or benzophenone-containing molecules. Here, we report a unique method for photo-crosslinking, employing the intrinsic photochemistry of the isoxazole, a common heterocycle in medicinal chemistry, to offer an alternative to existing strategies using more perturbing, extrinsic crosslinkers. In this initial report, this technique is applied both *in vitro* and *ex vivo*, used in a variety of common chemoproteomic workflows, and validated across multiple proteins, demonstrating the utility of isoxazole photo-crosslinking in a wide range of biologically relevant experiments.

---

Photo-crosslinking or photoaffinity labeling (PAL) connects small molecule drug discovery to proteomics, providing detailed information like ligand binding sites or global information like off-target profiling.^1^ Despite wide usage of photocrosslinking in chemical biology, few advancements have been made regarding the photochemistry employed. Emphasis has been on time-tested photoreactive groups such as aryl azides, diazirines, and benzophenones.^2–4^ However, the synthesis and attachment of these motifs is not always straightforward and may alter binding of the parent molecule. Work by the Woo and Parker laboratories has investigated how these existing methods function and how selectivity and efficiency can be optimized.^5–7^ A study from GlaxoSmithKline also enforced the utility of photo-crosslinkers in drug discovery, using diazirines in fragment-based screening to identify binders of protein targets.^8^ One example of an expansion of the photo-crosslinking toolbox has been the development of 2-aryl-5-carboxytetrazole (ACT).^9^ Upon irradiation, ACT is proposed to generate a highly reactive diaryl nitrile imine, which can undergo nucleophilic attack by nearby amino acid sidechains. Considering all of this, we posited that a novel photo-crosslinker would be a welcome addition to this underdeveloped area of chemical biology.

Isoxazoles represent a common motif in medicinal chemistry.^10^ Intriguingly, several studies have highlighted the labile nature of the isoxazole N-O bond. The Khlebnikov group has achieved transformations such as: rearrangement of vinyl-isoxazoles to pyrroles^11^, isoxazole-azirine and isoxazole-oxazole isomerization^12^, and azirine-oxazole conversion followed by reductive isoxazole ring-opening.^13^ The majority of these reactions utilized a catalyst. However, there is significant precedent for light-activated ring opening of isoxazoles.^14^ The isoxazole accesses high-energy intermediates reminiscent of photo-crosslinking motifs during these transformations, including nitrenes and azirines (**Scheme 1**).^15^ Indeed, muscimol and thiomuscimol, isoxazole-containing substrates of the gamma amino butyric acid receptor, have been shown to crosslink without exogenous photo-crosslinking groups, presumably due to the photoreactivity of the isoxazole.^16–18^ Isoxazoles represent an untapped functional group in terms of their innate ability to act as a minimalist photo-crosslinker and their presence in a wide variety of molecules in medicinal chemistry. We therefore sought to establish proof-of-principle for isoxazoles as photo-crosslinkers/PAL probes by demonstrating their utility in binding site validation and off-target analysis with implications for drug repurposing.

**Scheme 1.**
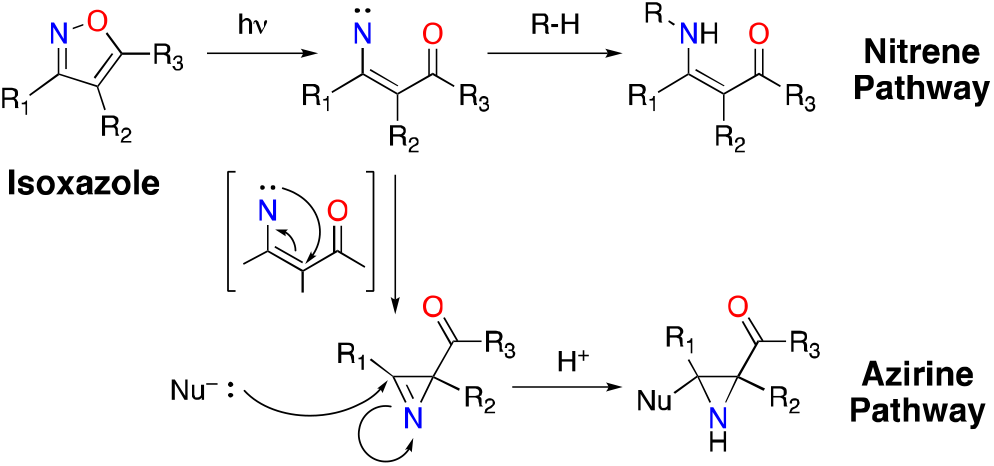
Isoxazole photo-crosslinking mechanisms.

α-Synuclein (αS) is a 14.5 kDa protein that is implicated in several neurodegenerative diseases. αS insoluble aggregates (fibrils) are highly concentrated in Lewy Bodies, the pathological hallmark of Parkinson’s disease.^19^ Initial photo-crosslinking studies were attempted using a pharmacophore previously identified in our efforts to develop positron emission tomography (PET) imaging probes for Parkinson’s disease. Isoxazole-containing molecules have been shown to bind to *in vitro* generated αS pre-formed fibrils (PFFs) with a nanomolar affinity.^20^ We envisioned a minimalist photo-crosslinker, with the core based around a light-reactive isoxazole fragment. Electronic effects on isoxazole crosslinking were assessed via attachment of electron-rich or electron-poor rings at the 3-position of the isoxazole. Inclusion of a propargyl ether enables subsequent experiments involving copper-catalyzed 3+2 cycloaddition (click) chemistry functionalization.^21–22^ Separation of the photoaffinity motif via decoupling of the photo-crosslinker and click chemistry handle has a two-part advantage: i) allowing the synthetic chemist more freedom in incorporating a simple alkyne tag and ii) reducing perturbation of binding by a larger photoaffinity handle.

The initial series of photo-crosslinkers (**AS1-7**) was validated against PFFs via matrix-assisted laser desorption ionization time-of-flight (MALDI-TOF) mass spectrometry (MS) and fluorescent SDS-PAGE analysis. Upon irradiation with 254 nm light, all probes formed covalent adducts with αS that retained the ability to be labeled via copper-catalyzed click chemistry using an Alexa Fluor 488 (AF488) azide probe. (**Figure 1B**) This shows that isoxazole crosslinking is not significantly affected by the electronic nature of the pendant ring system. Alternative wavelengths (365 nm, 427 nm) were examined as well, but resulted in decreased labeling efficiency. (**Supporting Information or SI, Figures S1-S2**) A mass adduct corresponding to isobaric labeling was noted via whole protein MALDI-TOF MS, indicating that the small molecule undergoes minimal photo-degradation after crosslinking despite the use of 254 nm light. (**Figure 1D**) An additional peak was observed via MALDI-TOF MS for all compounds, possibly due to in-source decay.

**Figure 1.**
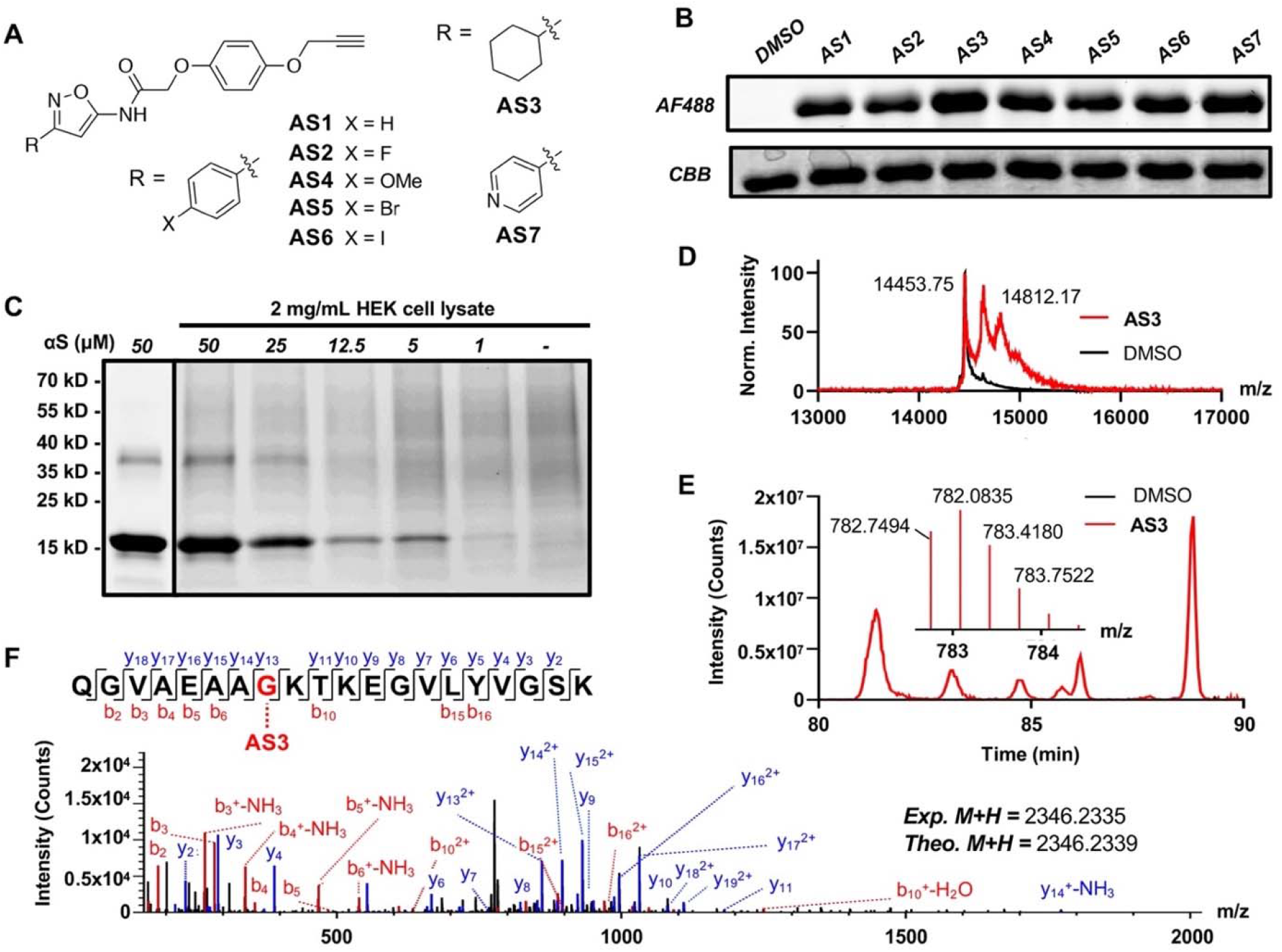
A. **AS1-7** isoxazole probes. B. Fluorescent SDS-PAGE of isoxazole-containing photo-crosslinkers irradiated at 254 nm in the presence of fibrillar αS. AF488: Fluorescent detection of αS photoproducts click labeled with AF488. CBB: Coomassie staining showing protein present in each lane. C. Crosslinking of fibrillar αS to **AS7** in the presence and absence of HEK cell cytosolic lysate. D. Whole protein MALDI-TOF MS of **AS3** crosslinking to fibrillar αS. E. αS fragment crosslinked to **AS3** identified via Orbitrap LC-MS/MS (extracted and parent ion chromatograms.) DMSO controls show spectra for un-crosslinked αS. F. Fragment (MS/MS) spectrum of identified **AS3** crosslinked peptide with annotated b and y ion series.

To demonstrate that our molecules enable target enrichment, cross-linked αS was labeled with desthiobiotin azide (DTBA), disaggregated by boiling, bound to streptavidin beads, and eluted with excess biotin. A product with the additional mass of DTBA was observed in the elution fraction, indicating successful enrichment. (**SI, Figure S4**) Probe **AS7** was examined in more detail for the ability to label αS in the presence of cytosolic human embryonic kidney (HEK) cell lysate. 100 μM **AS7** in the presence of 2 mg/ml HEK cell lysate showed a subtle decrease in αS labeling efficiency. This is likely attributable to non-specific binding of these hydrophobic probes, as has been previously observed in our laboratory.^20^ However, both samples exhibited a similar αS detection limit (1-5 μM), demonstrating the utility of the isoxazole as a novel photo-labeling motif even in the complex milieu of cell lysates. (**Figure 1C**)

Given the efficiency of *in vitro* photo-crosslinking, we wished to determine whether we could obtain binding site information for our probe molecules by performing bottom-up proteomics. αS contains a variety of small-molecule binding sites,^23^ and it is not obvious which of these are targeted by a given ligand. Upon irradiation of **AS3**, a substantial adduct was noted by whole-protein MALDI-TOF MS (**Figure 1D**). Upon tryptic digest, Orbitrap liquid chromatograph MS/MS (LC-MS/MS) was used to identify a match for residues 24-43 of αS (**Figure 1E**). Analysis of the fragment MS/MS spectrum allowed us to confirm the identity of this crosslinked peptide and localize the crosslink to Gly_31_ (**Figure 1F**). The majority of the other identified crosslinked peptides were also localized to this region (**SI, Figure S5**), consistent with previous mapping of the binding site of a similar molecule to this site using a diazirine.^20^ These results demonstrate the utility of isoxazole as a minimal photo-crosslinking motif in protein binding site analysis.

To validate the use of isoxazole photo-crosslinkers more broadly, we shifted focus to FDA-approved drugs, where at least 15 compounds are currently or have previously been approved. Amongst these molecules, sulfisoxazole (**SFX, Figure 2A**) is an antibacterial agent that functions as a competitive inhibitor of dihydropteroate synthase, which is necessary to bacterial folate biosynthesis.^24^ In addition to its role as an antibiotic, **SFX** has been identified as a hit in anticancer drug repurposing screens, where it is proposed to inhibit extracellular vesicle (exosome) release via interactions with endothelin receptor A (ETA).^25^ There has been some debate as to the mechanism of reduction in tumor burden,^26–27^ motivating **SFX** PAL studies. Regardless of previously identified targets, novel binders of **SFX** would be useful for drug-repurposing as well as benchmarking of isoxazole photochemistry.

**Figure 2.**
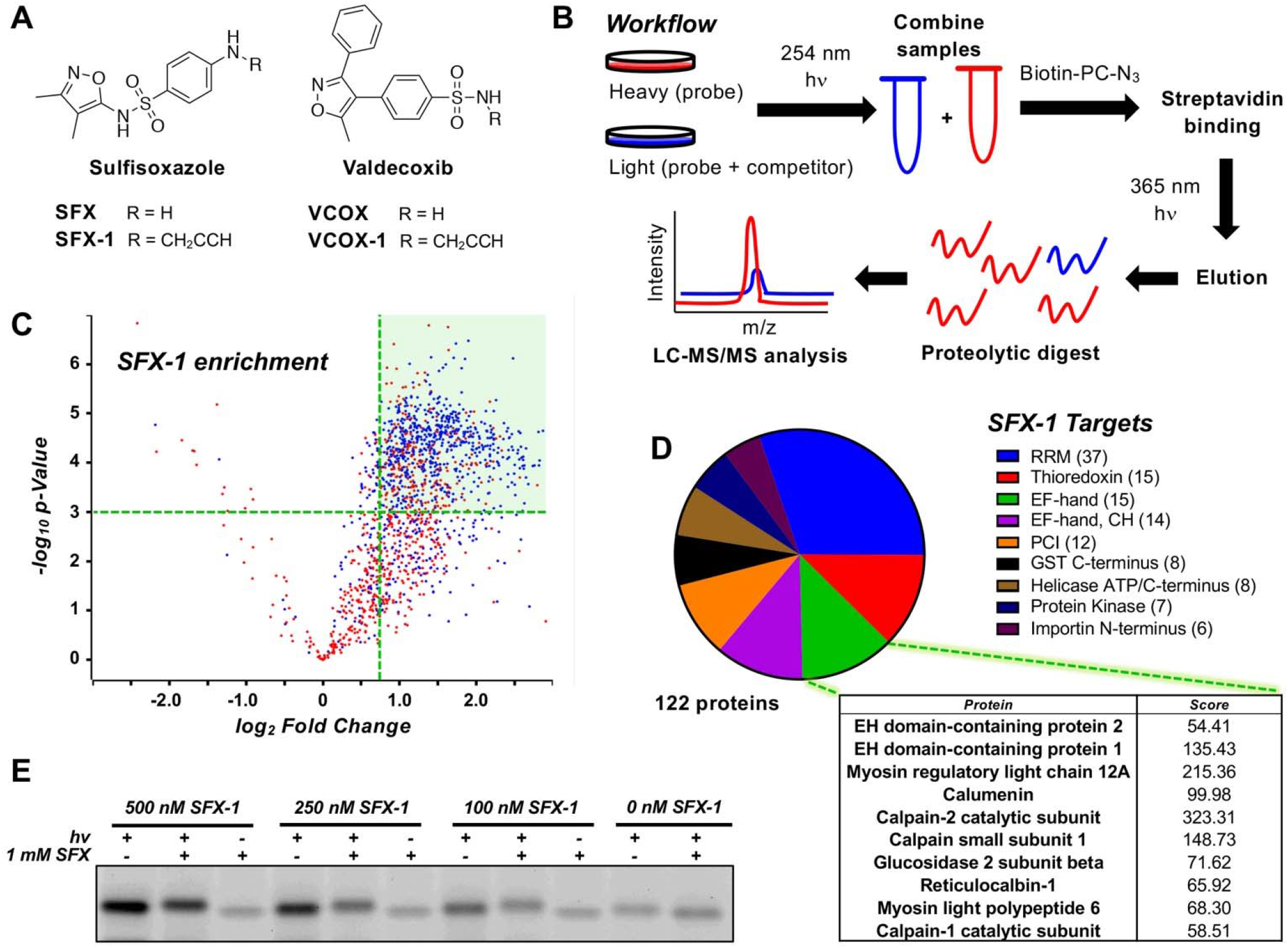
A. Structures of sulfisoxazole and valdecoxib probes. B. Schematic representation of the chemoproteomic workflow utilized in this study. C. Volcano plot representing **SFX-1** enrichment of proteins with an andromeda score greater than 15 colored in blue and the enrichment region for the top 10 EF-hand containing proteins highlighted in green. D. Pie chart representing proteins targeted by **SFX-1**, organized by k-means clustering of specific protein domains, with sections sized depending on number of proteins. See SI for details. E. Fluorescence SDS-PAGE analysis of CaM *in vitro* photo-crosslinking with **SFX-1**. Note: CaM stains weakly in the absence of **SFX**/**SFX-1**.

A sulfisoxazole probe (**SFX-1, Figure 2A**) was synthesized to contain an alkyne for subsequent labeling and enrichment studies. **SFX-1** was first validated by photo-crosslinking in the presence of 2 mg/mL HEK cell lysate with subsequent conjugation to tetramethyl rhodamine (TAMRA) azide. From this, the optimal irradiation time was determined to be ~10 min, as a time-dependent decrease in protein banding upon prolonged UV exposure was observed. (**SI, Figure S6**) The application of this probe in a chemoproteomic workflow was then examined quantitatively in cytosolic MDA-MB-231 cell lysate via stable isotopic labeling using amino acids in cell culture (SILAC).^28^ The SILAC lysates were treated with either 250 μM probe (heavy “H” samples, **SFX-1**) or 250 μM probe and 1 mM competitor (light “L” samples, **SFX-1** + **SFX**). The resulting samples were then crosslinked, click labeled with a photo-cleavable biotin azide, enriched on streptavidin beads, eluted via irradiation at 365 nm, and analyzed via LC-MS/MS. (**Figure 2B**)

A robust list of proteins was identified after filtering of contaminants (1555 total) that was further enriched (785 total) via thresholding (H/L ≥ 1.5, Andromeda score > 15, p-value < 0.05). (**Figure 2C**) Given the remaining complexity of the target list, a bioinformatic similarity-based approach was adopted to select potential high-value targets of **SFX**. Proteins were analyzed via domain annotation in Uniprot (60% annotated), and the proteins pertaining to the top-10 enriched domains (122 total) were one-hot encoded and grouped using k-means clustering. (**Figure 2D**)

The top enriched domain was determined to be the RNA recognition motif (RRM) followed by the EF-hand. Both domain families represent interesting targets. The RRM domain protein list was mainly composed of heterogenous nuclear ribonucleoproteins, which have been shown to be enriched and shuttled in extracellular vesicles.^29^ The EF-hand domain is a simple helix-loop-helix motif found in a suite of calcium-binding proteins and is also a high-value target as it may have implications in exosome biogenesis, the proposed mechanism of **SFX** as an anti-cancer agent.^30^ Aside from the EF-hand domain proteins, a number of other calcium-modulating proteins were enriched that have shown ties to exosome formation including: synaptotagmin^31^, calpain^32^, and the annexin family of proteins^33–34^.

As a secondary cross-validation of **SFX-1** results, photo-crosslinking was conducted in parallel with a structurally similar drug molecule with different pharmacological properties. Valdecoxib (**VCOX**, **Figure 2A**) is a cyclooxygenase 2 (COX2) inhibitor that was FDA-approved until 2004 when it was withdrawn due to side-effects.^35^ An alkyne-containing derivative was synthesized (**VCOX-1**) and photo-crosslinking SILAC studies were undertaken in parallel with **SFX-1**. Lysates were treated with either 500 μM probe (“H” samples, **VCOX-1**) or 500 μM probe and 500 μM competitor (“L” samples, **VCOX1** + **VCOX**). Samples were then crosslinked, enriched as in sulfisoxazole SILAC studies, and analyzed by LC-MS/MS. Although overlapping targets were identified between **SFX-1** and **VCOX-1**, the drugs showed differences in their enrichment profiles. Filtering **SFX-1** protein targets by removing proteins enriched in **VCOX-1** treated samples (H/L <= 1.5 removed) yielded a list containing several EF-hand proteins, similar to what was identified in the **SFX**/**SFX-1** experiment by domain clustering. This evidence further confirms labeling of the EF-hand motif by **SFX-1**. (**SI, Figures S8-S9**)

To validate the observed trends, the EF-hand-containing **SFX** hit calmodulin (CaM) was expressed and crosslinking was performed *in vitro*. CaM showed photo-crosslinking with **SFX-1** and competition with various dosages of **SFX**. (**Figure 2E**) It should be noted that the SILAC experiments focused exclusively on cytosolic targets. While crosslinking to a protein with a molecular weight consistent with ETA was observed in membrane fractions (**SI, Figure S7**), we restricted our SILAC experiments to the cytosol (Triton-soluble fraction) to enable optimal enrichment on streptavidin beads without the risk of denaturing the proteome. The discovery of novel targets for **SFX** reinforces the ability of isoxazole photo-crosslinking methods to spearhead drug-repurposing screens using existing molecules from medicinal chemistry libraries and provides a rich data set for subsequent biological analysis.

In this initial report, we have demonstrated that isoxazole compounds can be activated with UV light for photocrosslinking to proteins and that this is compatible with most chemoproteomic workflows, including photo-cleavable biotin tags, with the addition of a bio-orthogonal reaction handle elsewhere on the molecule. We have validated isoxazole crosslinking in αS PFF experiments and showed that we identify the same binding site previously observed with a conventional diazirine tag. We have also provided insight as to the anti-cancer mechanism and interactome of **SFX**. The usefulness of isoxazoles as a photochemical motif emphasizes the need for the identification and development of additional novel photocrosslinkers, further enabling chemists to utilize functional, minimalist, and easily accessible small molecule probes. Indeed, studies of the photo-reactivity of other common medicinal chemistry heterocycles is already underway in our laboratory.

## Supporting information

SFX_VCOX_Comparisons

SFX_Proteomics_Data

## ASSOCIATED CONTENT

Supporting Information.

## AUTHOR INFORMATION

### Author Contributions

All experiments were conceived by M.G.L. and E.J.P. Chemical synthesis was performed by V.V.P. and M.G.L. All crosslinking and MALDI MS experiments were performed by M.G.L with assistance by S.X.P. Orbitrap MS experiments were performed by H.J.K. with data analysis by M.G.L. V.V.P. is supported by E. J. P. and R.M.H. H.J.K. is supported by B.A.G. The manuscript was written by M.G.L. and E.J.P. with contributions of all authors. All authors have given approval to the final version of the manuscript.

### Notes

The authors declare no competing financial interests.

## ACKNOWLEDGMENT

This research was supported by the National Institutes of Health (NIH R01-NS103873 to E.J.P., NIH U19-NS110456 to R.H.M. and E.J.P., and P01-CA196539 and R01-AI118891 to B.A.G.). Instruments supported by the NIH and National Science Foundation include NMR (NSF CHE-1827457) and mass spectrometers (NIH RR-023444 and NSF MRI-0820996) M.G.L. was supported by an Age Related Neurodegenerative Disease Training Grant fellowship (NIH T32-AG000255). H.J.K. was supported by an NIH Predoctoral Fellowship (NIH F31-AG069390). S.X.P. thanks the Vagelos Molecular Life Science Scholars program for support. M.G.L thanks Chloe M. Jones for recombinant calmodulin used during *in vitro* crosslinking studies.

