## Supporting Information for "Harnessing the Intrinsic Photochemistry of Isoxazoles for the Development of Chemoproteomic Crosslinking Methods"

**Table of Contents:**

*General Information* .......... **S2**
*Synthetic Chemistry Methods* .......... **S3**
*Protein Expression and Purification* .......... **S11**
*Photo-reactivity Studies* .......... **S14**
*Mass Spectrometry Analysis of α-Synuclein Crosslinking* .......... **S16**
*Sulfisoxazole and Valdecoxib Crosslinking* .......... **S17**
*MS/MS Data Acquisition and Analysis* .......... **S19**
*Figures* .......... **S22**
*References* .......... **S29**

**Associated Files:** Data sets for all proteomic experiments are supplied as separate Microsoft Excel spreadsheets in either **SFX_Proteomic_Data.xlsx** (**SFX-1**-only experiments) or **SFX_VCOX_Comparisons.xlsx** (**SFX/VCOX** comparison study). Python scripts utilized for data analysis are deposited on GitHub (https://github.com/ejp-lab/EJPLab_Computational_Projects/tree/master/PhotoCrosslinking).

***General Information***

All cell culture was conducted in a HEPA-filtered cell culture hood under sterile conditions. Buffers were made with MilliQ filtered (18 MΩ) water (Millipore; Billerica, MA, USA). DC protein assay kits were purchased from Bio-Rad (Bio-Rad; Hercules, CA, USA). Matrix-assisted laser desorption/ionization (MALDI) mass spectrometry (MS) data were collected with a Bruker Ultraflex III MALDI-TOF/TOF mass spectrometer (Billerica, MA, USA). UV/Vis absorbance spectra were obtained with a Thermo Fisher Scientific GENESYS 150 UV-Vis spectrophotometer (Thermo Fisher Scientific; Waltham, MA, USA). DC assay protein quantitation was conducted using a Tecan Spark microplate reader (Tecan; Mannedorf, Switzerland). SDS-PAGE gels were run using a Bio-Rad PowerPac Basic Power Supply. Gel images were obtained with a Bio-Rad ChemiDoc MP instrument. Protein constructs were purified on an Äkta pure FPLC (Cytiva; Marlborough, MA, USA). NMR spectra were obtained on a Bruker NEO 400 spectrometer. Electrospray ionization (ESI) mass spectra were obtained on a Waters Acquity Ultra Performance LC connected to a single quadrupole detector (SQD) mass spectrometer (Waters Corp.; Milford, MA, USA). High resolution electrospray ionization mass spectra (ESI-HRMS) were obtained on a Waters LCT Premier XE liquid chromatograph/mass spectrometer. Orbitrap liquid chromatrography MS/MS (LC-MS/MS) data were acquired on a Thermo Fisher Scientific QE-HF instrument. (Thermo Fisher Scientific; Waltham, MA, USA). Python version 3.7.5 was used for data anlaysis scripts. MaxQuant (version 1.6.17.0) was downloaded from https://www.maxquant.org/download_asset/maxquant/latest. ProteomeDiscoverer version used was 2.4. GraphPad Prism version used was 9.0.2.

*Synthetic Chemistry Methods*

**Chemical Reagents and Instruments**. Chemicals were obtained from commercial sources and used without further purification. Solvents were purchased from commercial sources and used as received unless stated otherwise. Reactions were performed at room temperature unless stated otherwise. Reactions were monitored by thin layer chromatography (TLC) on pre-coated silica 60 F254 aluminum plates (MilliporeSigma, Burlington, MA, USA), spots were visualized by ultraviolet (UV) light. Evaporation of solvents was performed under reduced pressure at 40 °C using a rotary evaporator. Flash column chromatography was performed on a Biotage® (Charlotte, NC, USA) Isolera One system equipped with Biotage® SNAP KP-Sil cartridges. Nuclear magnetic resonance (NMR) spectroscopy was performed on a Bruker (Billerica, MA, USA) Avance Neo 400 (400.17 MHz for $^1$H and 100.63 MHz for $^{13}$C) with chemical shifts (δ) reported in parts per million (ppm) relative to the solvent ($CDCl_3$, $^1$H 7.26 ppm, $^{13}$C 77.16 ppm; dimethyl sulfoxide (DMSO)-*d6*, $^1$H 2.50 ppm, $^{13}$C 39.52 ppm). Low resolution Liquid Chromatography Mass Spectrometry (LCMS) was carried out using a Waters (Milford, MA, USA) SQD equipped with an Acquity UPLC instrument in positive ion mode. High Resolution Mass Spectrometry (HRMS) for small molecules were obtained on a Waters LCT Premier XE LC/MS system.

**Scheme S1.** Synthesis of **AS1-7** compounds

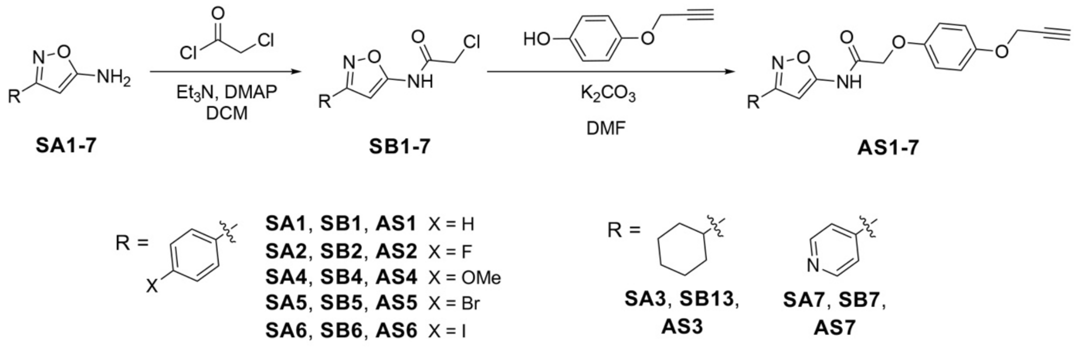


**General Procedure A.** An aryl isoxazol-5-amine (**SA1-7**, 0.40 mmol, 1.0 equiv.) and 4-dimethylamino-pyridine (DMAP, 5 mol%) were charged to an oven-dried round bottom flask with a stir bar. The flask was sealed with a rubber septum and sparged with argon. Dry $CH_2Cl_2$ (3.0

mL) and trimethylamine (3.0 equiv.) were then added to the flask, and the solution was cooled on ice for approximately 5 min. 2-chloroacetyl chloride (3 equiv.) was then added dropwise. The reaction was allowed to stir for 20 min before the ice was removed and the reaction was stirred overnight at 32 °C. The reaction was quenched with water (10 mL) and the reaction mixture was further diluted with water (30 mL) and $CH_2Cl_2$ (40 mL) and then layers were separated. The aqueous layer was further extracted 2 times with $CH_2Cl_2$ (40 mL). The combined organic layer was washed with saturated aqueous $NaHCO_3$ (50 mL) and saturated brine solution (50 mL), the organic layer was dried over $Na_2SO_4$ and concentrated under vacuum. The crude compound, **SB1-7**, was purified by flash column chromatography (gradient of 30-50% EtOAc/hexanes) to obtain the desired product in 60-80% yield.

**General Procedure B.** 4-(prop-2-yn-1-yloxy)phenol (0.35 mmol, 2.0 equiv.) and $K_2CO_3$ (3.0 equiv.) were charged to an oven-dried round bottom flask with a stir bar. The flask was then sealed with a rubber septum and sparged with argon and dry DMF (3.0 mL) was added to flask. The respective acetyl chloride **SB1-7** (1.0 equiv.) was solubilized in DMF (1 mL) and added slowly to the solution. The flask was then heated at 65 °C overnight. The reaction mixture was diluted with water (50 mL) and extracted with EtOAc (2x 50 mL). The combined organic layer was washed with saturated brine solution (50 mL), the organic layer was dried over $Na_2SO_4$ and concentrated under vacuum. The crude compound, **AS1-7**, was purified by flash column chromatography (gradient of 40-70% EtOAC/hexanes) to obtain desired product in 60-80% yield.

***N*-(3-phenylisoxazol-5-yl)-2-(4-(prop-2-yn-1-yloxy)phenoxy)acetamide (AS1)**

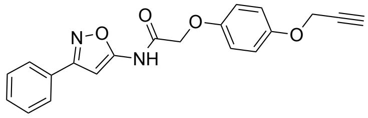


TLC (Hexanes:EtOAc, 65:35 v/v): $R_f$ = 0.35; Off White Solid; yield 74%; **$^1$H NMR (600 MHz, $CDCl_3$):** δ 9.20 (s, 1H), 7.82 (dd, *J* = 7.4, 2.2 Hz, 2H), 7.46 (d, *J* = 1.0 Hz, 1H), 7.45 (d, *J* = 2.4 Hz, 2H), 6.98 (d, *J* = 9.2 Hz, 2H), 6.94 (d, *J* = 9.2 Hz, 2H), 6.82 (s, 1H), 4.66 (d, *J* = 2.4 Hz, 2H), 4.65 (s, 2H), 2.52 (t, *J* = 2.4 Hz, 1H); **$^{13}$C NMR (150 MHz, CDCl3):** δ 163.7, 162.8, 158.5, 152.1,

150.3, 129.2, 127.9, 127.8, 125.8, 115.4, 114.9, 86.6, 77.4, 74.6, 66.9, 55.4; **HRMS-(ESI-TOF) (*m/z*):** calcd for $C_{20}H_{16}N_2O_4$ [M+Na]$^+$ 371.1008; found 371.1008.

***N*-(3-(4-fluorophenyl)isoxazol-5-yl)-2-(4-(prop-2-yn-1-yloxy)phenoxy)acetamide (AS2)**

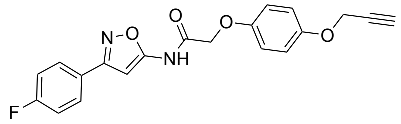


TLC (Hexanes:EtOAc, 70:30 v/v): $R_f$ = 0.25; Off White Solid; yield 52%; **$^1$H NMR (600 MHz, $CDCl_3$):** δ 9.18 (s, 1H), 7.82 (dd, *J* = 8.4, 5.3 Hz, 2H), 7.15 (t, *J* = 8.6 Hz, 2H), 6.98 (d, *J* = 9.2 Hz, 2H), 6.94 (d, *J* = 9.2Hz, 2H), 6.78 (s, 1H), 4.67 (d, *J* = 2.4 Hz, 2H), 4.65 (s, 2H), 2.52 (t, *J* = 2.4 Hz, 1H); **$^{13}$C NMR (150 MHz, CDCl3):** δ 164.8, 164.7, 163.1, 162.9, 159.6, 152.3 (d, *J* = 276.9 Hz), 128.8 (d, *J* = 8.2 Hz), 125.1 (d, *J* = 3.2 Hz), 116.4, 116.0 (d, *J* = 22.0 Hz), 115.9, 87.5, 78.5, 75.6, 67.9, 56.4; **HRMS-(ESI-TOF) (*m/z*):** calcd for $C_{20}H_{15}FN_2O_4$ [M+H]$^+$ 367.1094; found 367.1094.

***N*-(3-cyclohexylisoxazol-5-yl)-2-(4-(prop-2-yn-1-yloxy)phenoxy)acetamide (AS3)**

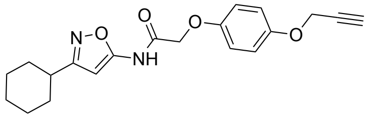


TLC (Hexanes:EtOAc, 65:35 v/v): $R_f$ = 0.30; White Solid; yield 57%; **$^1$H NMR (600 MHz, $CDCl_3$):** δ 9.06 (s, 1H), 6.96 (d, *J* = 9.1 Hz, 2H), 6.91 (d, *J* = 9.1Hz, 2H), 6.33 (s, 1H), 4.66 (d, *J* = 2.4 Hz, 2H), 4.60 (s, 2H), 2.72-2.69 (m, 1H), 2.51 (t, *J* = 2.4 Hz, 1H), 1.98-1.95 (m, 2H), 1.83-1.80 (m, 2H), 1.74-1.71 (m, 1H), 1.50-1.43 (m, 2H), 1.41-1.34 (m, 2H), 1.29-1.25 (m, 1H); **$^{13}$C NMR (150 MHz, CDCl3):** δ 170.2, 164.6, 158.5, 153.1, 151.4, 116.4, 115.9, 87.9, 78.5, 75.6, 67.8, 56.4, 36.3, 31.8, 25.9, 25.8; **HRMS-(ESI-TOF) (*m/z*):** calcd for $C_{20}H_{22}N_2O_4$ [M+H]$^+$ 355.1658; found 355.1657.

***N*-(3-(4-methoxyphenyl)isoxazol-5-yl)-2-(4-(prop-2-yn-1-yloxy)phenoxy)acetamide (AS4)**

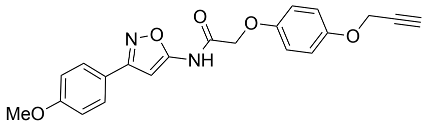

TLC (Hexanes:EtOAc, 60:40 v/v): $R_f$ = 0.30; Off White Solid; yield 68%; **$^{1}$H NMR (600 MHz, $CDCl_3$):** δ 9.15 (s, 1H), 7.76 (d, *J* = 8.8 Hz, 2H), 6.99-6.96 (m, 4H), 6.94 (d, *J* = 9.2 Hz, 2H), 6.76 (s, 1H), 4.66 (d, *J* = 2.4 Hz, 2H), 4.64 (s, 2H), 3.85 (s, 3H), 2.52 (t, *J* = 2.4 Hz, 1H); **$^{13}$C NMR (150 MHz, CDCl3):** δ 164.7, 163.4, 161.2, 159.2, 153.1, 151.4, 128.2, 121.4, 116.4, 115.8, 114.3, 87.4, 78.5, 75.6, 67.9, 56.4, 55.4; **HRMS-(ESI-TOF) (*m/z*):** calcd for $C_{21}H_{18}N_2O_5$ $[M+H]^+$ 379.1294; found 379.1291.

***N*-(3-(4-bromophenyl)isoxazol-5-yl)-2-(4-(prop-2-yn-1-yloxy)phenoxy)acetamide (AS5)**

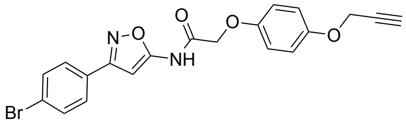

TLC (Hexanes:EtOAc, 70:30 v/v): $R_f$ = 0.25; White Solid; yield 55%; **$^{1}$H NMR (600 MHz, $CDCl_3$):** δ 9.17 (s, 1H), 7.70 (d, *J* = 8.4 Hz, 2H), 7.60 (d, *J* = 8.4 Hz, 2H), 6.98 (d, *J* = 9.1Hz, 2H), 6.94 (d, *J* = 9.1Hz, 2H), 6.79 (s, 1H), 4.67 (d, *J* = 2.2 Hz, 2H), 4.65 (s, 2H), 2.52 (t, *J* = 2.2 Hz, 1H); **$^{13}$C NMR (150 MHz, CDCl3):** δ 164.7, 162.9, 159.7, 153.2, 151.3, 132.2, 128.3, 127.4, 124.6, 116.4, 115.9, 87.4, 78.5, 75.6, 67.9, 56.4; **HRMS-(ESI-TOF) (*m/z*):** calcd for $C_{20}H_{15}BrN_2O_4$ $[M+H]^+$ 427.0293; found 427.0291.

***N*-(3-(4-iodophenyl)isoxazol-5-yl)-2-(4-(prop-2-yn-1-yloxy)phenoxy)acetamide (AS6)**

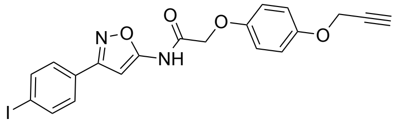

TLC (Hexanes:EtOAc, 70:30 v/v): $R_f$ = 0.25; Off White Solid; yield 51%; **$^{1}$H NMR (600 MHz, $CDCl_3$):** δ 9.18 (s, 1H), 7.80 (d, *J* = 8.4 Hz, 2H), 7.56 (d, *J* = 8.4 Hz, 2H), 6.98 (d, *J* = 9.1Hz, 2H), 6.94 (d, *J* = 9.1Hz, 2H), 6.79 (s, 1H), 4.67 (d, *J* = 2.2 Hz, 2H), 4.65 (s, 2H), 2.52 (t, *J* = 2.3 Hz, 1H); **$^{13}$C NMR (150 MHz, CDCl3):** δ 164.7, 163.0, 159.7, 153.2, 151.3, 138.1, 128.4, 128.3,

116.4, 115.9, 96.5, 87.4, 78.5, 75.7, 67.9, 56.4; **HRMS-(ESI-TOF) (*m/z*):** calcd for $C_{20}H_{15}IN_2O_4$ $[M+H]^+$ 475.0155; found 475.0147.

**2-(4-(prop-2-yn-1-yloxy)phenoxy)-*N*-(3-(pyridin-4-yl)isoxazol-5-yl)acetamide (AS7)**

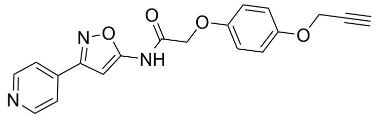

TLC (Hexanes:EtOAc, 35:65 v/v): $R_f$ = 0.35; Light brown Solid; yield 45%; **$^1$H NMR (600 MHz, DMSO-d$_6$):** δ 12.0 (s, 1H), 8.72 (d, *J* = 5.4 Hz, 2H), 7.85 (d, *J* = 6.0 Hz, 2H), 6.95 (s, 4H), 9.93 (s, 1H), 4.78 (s, 2H), 4.73 (d, *J* = 2.3 Hz, 2H), 3.53 (t, *J* = 2.3 Hz, 1H); **$^{13}$C NMR (150 MHz, DMSO-d$_6$):** δ 166.5, 163.0, 161.5, 152.7, 152.2, 151.0, 136.4, 121.2, 116.3, 116.0, 87.2, 79.9, 78.5, 67.5, 56.3; **HRMS-(ESI-TOF) (*m/z*):** calcd for $C_{19}H_{15}N_3O_4$ $[M+H]^+$ 350.1141; found 350.1126.

**4-(prop-2-yn-1-yloxy)phenol**

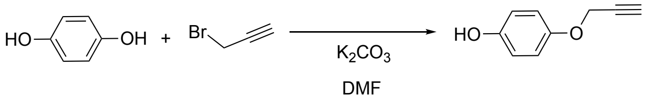


A flask of dihydroquinone (90.818 mmol, 3.0 eq.), DMF (60 ml), and $K_2CO_3$ (93.846 mmol, 3.1 eq.) under argon was charged with propargyl bromide (30.273 mmol, 1.0 eq.) dropwise with stirring. Immediately following addition, the flask was heated to 60 °C and the reaction was allowed to stir overnight. The crude reaction mixture was then filtered through Celite, and the filter cake was washed three times with EtOAc (25 ml). The resulting solution was then acidified with 1M HCl (200 ml) and extracted three times with EtOAc (50 ml). The organic phase was then washed twice with brine (200 ml), dried over $Na_2SO_4$, and adsorbed onto silica gel. The crude product was then purified by Biotage flash column chromatography (100 g column, gradient 0-40% EtOAc/hexanes, 30 column volumes) to obtain 4-(prop-2-yn-1-yloxy)phenol in 60% yield. TLC (Hexanes:EtOAc, 50:50 v/v): $R_f$ = 0.56; off-white solid; **$^1$H NMR (600 MHz, CDCl3):** δ 6.88 (d, J = 9.0 Hz, 2H), 6.78 (d, J = 9.0 Hz, 2H), 4.85 (s, 1H), 4.63 (d, J = 2.4 Hz, 2H), 2.51 (d, J

= 4.8 Hz, 1H); **$^{13}$C NMR (151 MHz, CDCl3):** δ 151.84, 150.31, 116.54, 116.20, 78.95, 75.55, 56.84; **LRMS-(ESI-TOF) (*m/z*):** calcd for $C_9H_8O_2$ $[M+H]^+$ 149.060; found 149.028.

***N*-(3,4-dimethylisoxazol-5-yl)-4-(prop-2-yn-1-ylamino)benzenesulfonamide (SFX-1)**

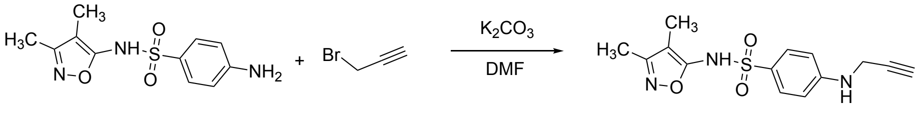


*N*-(3,4-dimethylisoxazol-5-yl)-4-(prop-2-yn-1-ylamino)benzenesulfonamide (**SFX**, 0.40 mmol, 1.0 equiv.) and $K_2CO_3$ (0.40 mmol, 0.75 equiv.) were charged to an oven-dried round bottom flask with a stir bar. The flask was sealed with a rubber septum and sparged with argon. Dry DMF (2.0 mL) was then added to flask, and the solution was cooled on ice for approximately 5 min. Propargyl bromide solution, 80 wt.% in toluene (0.40 mmol, 0.5 equiv.), was then added dropwise. The reaction was allowed to stir for 20 min before the ice was removed and the reaction was stirred for 5 h at room temperature. The reaction was quenched with water (25 mL) and ethyl acetate (25 mL) and then layers were separated. The aqueous layer was further extracted 2 times with ethyl acetate (25 mL). The combined organic layer was washed with saturated brine solution (50 mL), the organic layer was dried over $Na_2SO_4$ and concentrated under vacuum. The crude compound was purified by flash column chromatography (gradient of 30-70% EtOAC/hexanes) to obtain the desired product **SFX-1** in 48% yield. TLC (Hexanes:EtOAc, 50:50 v/v): $R_f$ = 0.35; Off White Solid; yield 48%; **$^1$H NMR (600 MHz, CDCl$_3$):** δ 7.52 (d, *J* = 8.7 Hz, 2H), 6.64 (d, *J* = 8.7 Hz, 2H), 4.25 (br, 2H), 4.23 (d, *J* = 2.5 Hz, 2H), 2.23 (s, 3H), 2.18 (t, *J* = 2.5 Hz, 1H), 1.97 (s, 3H); **$^{13}$C NMR (150 MHz, CDCl3):** δ 161.9, 157.4, 151.5, 130.1, 125.4, 114.0, 111.8, 76.6, 73.9, 39.6, 11.0, 7.0; **HRMS-(ESI-TOF) (*m/z*):** calcd for $C_{14}H_{15}N_3O_3S$ $[M+H]^+$ 305.0834; found 306.0897.

**4-(5-methyl-3-phenylisoxazol-4-yl)-*N*-(prop-2-yn-1-yl)benzenesulfonamide (VCOX-1)**

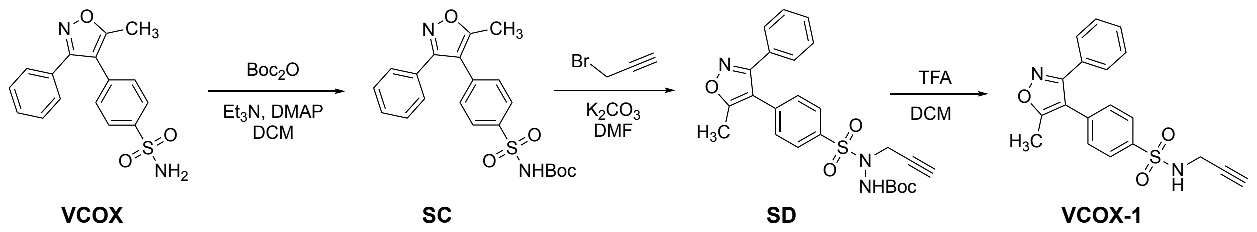



4-(5-methyl-3-phenylisoxazol-4-yl)benzenesulfonamide (**VCOX**, 0.40 mmol, 1.0 equiv.) and DMAP (2.5 mol%) were charged to an oven-dried round bottom flask with a stir bar. The flask was then sealed with a rubber septum, sparged with argon, and dry DCM (5.0 mL) was added to flask. Triethyl amine (0.40 mmol, 1.5 equiv.) was added and then di-*tert*-butyl dicarbonate (0.40 mmol, 1.1 equiv.) was solubilized in DCM (3 mL) and added slowly to the solution. The flask was then stirred at room temperature for 3h. The reaction was quenched with water (50 mL) and EtOAc (50 mL) and then layers were separated. The aqueous layer was further extracted with EtOAc (50 mL). The combined organic layer was washed with water (50 mL) and saturated brine solution (50 mL), then the organic layer was dried over $Na_2SO_4$ and concentrated under vacuum. The crude compound, *t*ert-butyl ((4-(5-methyl-3-phenylisoxazol-4-yl)phenyl)sulfonyl)carbamate (**SC),** was subjected to the next step without further purification.

**SC** (0.40 mmol, 1.0 equiv.) and $K_2CO_3$ (0.40 mmol, 2 equiv.) were charged to an oven-dried round bottom flask with a stir bar. The flask was sealed with a rubber septum and sparged with argon. Dry DMF (2.0 mL) was then added to flask, and a propargyl bromide solution, 80 wt.% in toluene (0.40 mmol, 1.1 equiv.), was then added dropwise. The reaction was stirred for 20 h at room temperature. The reaction was quenched with water (50 mL) and ethyl acetate (50 mL) and then layers were separated. The aqueous layer was further extracted 2 times with EtOAc (50 mL). The combined organic layer was washed 2 times with water (50 mL) and saturated brine solution (100 mL), then the organic layer was dried over $Na_2SO_4$ and concentrated under vacuum. The crude compound, *t*ert-butyl 2-((4-(5-methyl-3-phenylisoxazol-4-yl)phenyl)sulfonyl)-2-(prop-2-yn-1-yl)hydrazine-1-carboxylate (**SD**), was subjected to the next step without further purification.

**SD** (0.40 mmol, 1.0 equiv.) was charged to an oven-dried round bottom flask with a stir bar. The flask was then sealed with a rubber septum, sparged with argon, and dry DCM (5.0 mL) was added to flask. Trifluoroacetic acid (0.40 mmol, 5 equiv.) was added slowly to the solution at room temperature and reaction mixture stirred further for 3 h. The reaction mixture was concentrated under vacuum and the crude compound was dissolved in DCM (100 mL). A 1 M NaOH solution was added to adjust the pH to 7. Water was added and then layers were separated. The aqueous layer was further extracted with DCM (50 mL). The combined organic layer was washed with saturated brine solution (100 mL), then the organic layer was dried over $Na_2SO_4$ and concentrated under vacuum. The crude compound was purified by flash column chromatography (gradient of 40-70% EtOAC/hexanes) to obtain desired product **VCOX-1** in 68 % yield over 3 steps. TLC (DCM:MeOH, 95:05 v/v): $R_f$ = 0.40; White Solid; yield 68%; **$^1$H NMR (600 MHz, $CDCl_3$):** δ 7.88 (d, *J* = 8.4 Hz, 2H), 7.38 (t, *J* = 6.8 Hz, 3H), 7.32 (t, *J* = 8.3 Hz, 4H), 4.81 (t, *J* = 4.4 Hz, 1H), 3.90 (dd, *J* = 4.4, 2.5 Hz, 2H), 2.48 (s, 3H), 2.07 (t, *J* = 2.5 Hz, 1H); **$^{13}$C NMR (150 MHz, CDCl3):** δ 167.3, 161.1, 138.8, 135.6, 130.3, 129.7, 128.7, 128.6, 128.5, 127.8, 114.5, 77.9, 73.0, 32.9, 11.7; **HRMS-(ESI-TOF) (*m/z*):** calcd for $C_{19}H_{16}N_2O_3S$ $[M+H]^+$ 353.0960; found 353.0955.

***Protein Expression and Purification***

**α-Synuclein Production.** α-Synuclein (αS) was expressed, purified, and aggregated as previously described, with slight modifications.[1] Human αS with a C-terminal intein-$His_6$ fusion was transformed into *Escherichia coli* (*E. coli*) BL21 cells and plated on ampicillin plates (100 µg/ml). A single colony was then inoculated into a 5 ml primary culture containing ampicillin (100 µg/ml) in Luria-Bertain (LB) media and grown for 5-6 h with shaking (250 rpm) at 37 °C. The primary culture was then transferred to 1 L LB containing ampicillin (100 µg/ml) and grown as previously described until reaching an optical density ($OD_{600}$) of 0.8-1.0. At this stage, protein production was induced by the addition of isopropyl β-D-1-thiogalactopyranoside (IPTG) to 1 mM final concentration, and the culture was grown with shaking (250 rpm) overnight at 18 °C. The following day, cells were harvested by centrifugation at 4 °C for 20 min at 4000 rpm (Sorvall GS3 rotor). Pellets were then re-suspended in 20 ml/L culture of 40 mM Tris, pH 8.3 supplemented with EDTA-free protease inhibitor tablets (Pierce Biotechnology; Waltham, MA, USA) and transferred to a metal cup for sonication. Cells were lysed by sonication on ice with a Q700 probe sonicator (QSonica LLC; Newtown, CT, USA) with the following settings: Amplitude 50, Process Time 2-3 min, Pulse-ON Time 1 s, Pulse-OFF Time 1 s. Crude lysate was then transferred to 50 ml centrifugation tubes and clarified via centrifugation at 14,000 rpm for 45 min (Sorvall SS34 rotor). Following centrifugation, supernatant was removed and transferred to a 50 ml Falcon tube. 5 ml of nickel agarose resin (GoldBio; St. Louis, MO, USA) was added, and the lysate-nickel mixture was incubated with nutation at 4 °C for 1-2 h. Lysate-nickel mixture was then poured into a 20 mL fritted column, and the flowthrough was saved. The remaining resin was then washed with ~20 mL Wash Buffer 1 (50 mM HEPES buffer, pH 7.5), ~20 ml Wash Buffer 2 (50 mM HEPES, 5 mM imidazole, pH 7.5), and eluted with 12 mL Elution Buffer (50 mM HEPES, 300 mM imidazole, pH 7.5). 2-mercaptoethanol (Bio-Rad Laboratories; Hercules, CA, USA) was then added to crude lysate (200 mM final concentration), and the mixture was allowed to incubate with nutation at room temperature overnight. The resulting cleaved protein was then dialyzed against 20 mM Tris pH 8.0 for 8-10 h. The resulting dialysate was then treated with 5 mL nickel agarose resin (GoldBio) and incubated with nutation at 4 °C for 1-2 h. The mixture was then applied to a 20 mL fritted column and flowthrough containing αS was collected in a 15 ml Falcon tube. The resulting enriched protein mixture was then dialyzed against 20 mM Tris, pH 8.0 overnight and purified via FPLC using a 5 ml HiTrap Q-HP column (Cytiva; Marlborough, MA, USA) using the following method: Buffer A: 20 mM Tris, pH 8.0; Buffer B: 20 mM Tris, 1 M NaCl, pH 8.0; Gradient: 0% Buffer B – 5 column volumes, 0-10% Buffer B – 5 column volumes, 20-30% Buffer B – 20 column volumes, 30-100% Buffer B – 10 column volumes; flowrate 3 mL/min. The resulting fractions were then assessed for purity via MALDI-TOF MS, and pure fractions were combined. Protein was then concentrated, and buffer exchanged into phosphate-buffered saline (PBS, NaCl 0.137 M, KCl 0.0027 M, $Na_2HPO_4$ 0.01 M, $KH_2HPO_4$ 0.0018 M) to a final concentration of 100-200 µM via Amicon 3 kDa MWCO filters (Millipore Sigma; St. Louis, MO, USA). Purified protein was aliquoted into 1.5 mL tubes and stored at -80 °C until further use.

**Calmodulin Production.** Calmodulin (CaM) was prepared as described previously.[2] Briefly, CaM with a C-terminal intein-$His_6$ fusion was transformed into *E. coli* BL21 cells and plated on ampicillin plates (100 µg/mL). A single colony was then inoculated into a 5 ml primary culture containing ampicillin (100 µg/mL) in non-inducing media (NIM, prepare as described previously[3]) and grown for 5-6 h with shaking (250 rpm) at 37 °C. 12.5 ml of primary culture was then transferred to 250 ml secondary culture containing auto-inducing media (AIM, prepare as described previously[3]) supplemented with ampicillin (100 µg/mL) and grown at 37 °C with shaking at 250 rpm for four h. The temperature and shaking speed were then decreased to 30 °C and 200 rpm respectively, and culture was allowed to incubate for 18-22 h. The following day, cells were harvested by centrifugation at 4 °C for 20 min at 4000 rpm (Sorvall GS3 rotor). Pellets were then re-suspended in 20 mL/L culture of 40 mM Tris, pH 8.3 supplemented with EDTA-free protease inhibitor tablets (Pierce Biotechnology) and transferred to a metal cup for sonication. Cells were lysed by sonication on ice with a Q700 probe sonicator with the following settings: Amplitude 50, Process Time 2-3 min, Pulse-ON Time 1 s, Pulse-OFF Time 1 s. Crude lysate was then transferred to 50 ml centrifugation tubes and clarified via centrifugation at 14,000 rpm for 45 min (Sorvall SS34 rotor). Following centrifugation, supernatant was removed and transferred to a 50 mL Falcon tube. 5 ml of nickel agarose resin (GoldBio) was added, and the lysate-nickel mixture was incubated with nutation at 4 °C for 1-2 h. Lysate-nickel mixture was then poured into a 20 mL fritted column, and the flowthrough was saved. The remaining resin was then washed with ~20 mL Wash Buffer 1 (50 mM HEPES buffer, pH 7.5), ~20 ml Wash Buffer 2 (50 mM HEPES, 5 mM imidazole, pH 7.5), and eluted with 12 mL Elution Buffer (50 mM HEPES, 300 mM imidazole, pH 7.5). 2-mercaptoethanol (Bio-Rad Laboratories) was then added to crude lysate (200 mM final concentration), and the mixture was allowed to incubate with nutation at room temperature overnight. The resulting cleaved protein was then dialyzed against 20 mM Tris pH 8.0 for 8-10 h. The resulting dialysate was then treated with 5 mL nickel agarose resin (GoldBio) and incubated with nutation at 4 °C for 1-2 h. The mixture was then applied to a 20 mL fritted column and flowthrough containing CaM was collected in a 15 mL Falcon tube. The resulting enriched protein mixture was then dialyzed against 20 mM Tris, pH 8.0 overnight and purified via FPLC using a 5 ml HiTrap Q-HP column (Cytiva) using the following method: Buffer A: 20 mM Tris, pH 8.0; Buffer B: 20 mM Tris, 0.5 M NaCl, pH 8.0; Gradient: 0-100% Buffer B – 100 minutes; flowrate 3 mL/min. The resulting fractions were then assessed for purity via MALDI-TOF MS, and pure fractions were combined. Protein was then concentrated to a final concentration of 100-200 μM via Amicon 3 kDa MWCO filters (Millipore Sigma). Purified protein was aliquoted into 1.5 mL tubes and stored at -80 °C until further use.

**α-Synuclein Fibril Preparation.** Fibrils were prepared by first diluting monomeric αS to 100 μM in PBS in a 1.5 mL tube (500 μL final volume). Tubes were then sealed with both Teflon tape and parafilm and were shaken for 7 days at 37 °C in an Ika MS 3 control orbital shaker (Wilmington, NC, USA) at 1300 rpm.

### *Photo-reactivity Studies*

**Probe Photophysical Characterization.** All probes (**AS1-7**, **SFX**, **VCOX**) were diluted from a 10 mM DMSO stock to 100 μM final concentration in 50/50 MeOH/$H_2O$. The probe solution was then transferred to a quartz cuvette and analyzed from 210-400 nm using a UV-Vis spectrophotometer (Thermo Fisher Scientific; Waltham, MA) blanked against the diluent. Raw spectra were plotted in GraphPad Prism 9. These data are shown in **Figure S1.**

**Photo-crosslinking and Fluorescence SDS-PAGE Analysis.** Fibrils (50-100 μM) prepared as described above were treated with 100 μM **AS1-7b** probe molecule from a DMSO stock (50 μL total volume, 1% DMSO) in a 1.5 mL tube. The sample was incubated at 37 °C for 1 hour. Following incubation, the lid was opened, and the sample was irradiated with shaking (500-750 rpm) for 10 min, with the lamp (UVP Multiple Ray Lamp, 254 nm, 8 watt; Analytik Jena; Jena, Germany) positioned approximately 1-2 cm from the top of the tube. Following irradiation, fibrils were directly subjected to copper-catalyzed click chemistry. Briefly, 16.5 μL of PBS was added to each sample to pre-adjust for the final reaction volume. 1.0 μL of Atto488-azide (Millipore Sigma) or TAMRA-azide (Lumiprobe; Cockeysville, MD, USA) was then added to each sample. Click-mix was made by pre-mixing 1.875 μL $CuSO_4$ (25 mM) with 1.875 μL *tris*(3-hydroxypropyltriazolylmethyl)amine (THPTA, 50 mM) per reaction/sample (i.e. 10 samples = 18.75 μL of each). 3.75 μL of this Click-mix was then charged to each sample ($CuSO_4$ final concentration: 500 μM, THPTA final concentration: 1.25 mM). Finally, 3.75 μL of sodium ascorbate (40 mM) was added (final concentration: 2 mM). Reactions were allowed to react with gentle agitation at room temperature for 1.5-2 h. Reactions were then quenched via the addition 25 μL 4xLDS loading buffer (Thermo Fisher; Waltham, MA, USA) with dithiothreitol (DTT, 200 mM) and were boiled for 30 min to disaggregate fibrillar species. 20 μL of the resulting solution was then separated via SDS-PAGE and imaged for fluorescence on a Chemidoc gel imager (Bio-Rad).

**Optimal Wavelength and Irradiation Time Determination.** Fibrils were treated as described above with varying wavelengths of irradiation (UVP Multiple Ray Lamp, 254 nm, 365 nm, and 427 nm; 8 watt; Analytik Jena) for 10 min or at 254 nm, for varying amounts of time. Click chemistry with Atto488-azide and analysis via fluorescence SDS-PAGE were conducted as described above. Wavelength data are shown in **Figure S2.**

**Crosslinking Visualization via Enhanced Chemiluminescence.** αS fibrils were treated as described above with irradiation at 254 nm for varying amounts of time. Click chemistry with biotin azide (Click Chemistry Tools; Scottsdale, AZ, USA) was then conducted as described for fluorophore azides. Proteins were then transferred to 0.2 μm nitrocellulose paper via wet transfer using an XCell blot II module (Invitrogen) for 1.5 h, using 1xNuPage transfer buffer. The blot was then blocked overnight using 3% bovine serum albumin (BSA; Millipore Sigma; St. Louis, MO, USA) in PBS with 0.1% Tween-20 (PBST). The following day, the blot was treated with streptavidin-horse radish peroxidase (strep-HRP; Thermo Fisher; Waltham, MA, USA) at 1:50000 dilution of 1 mg/mL stock for 1 h. The sample was then washed 5 times with PBST for 5 min each and exposed to ECL substrate (Pierce Biotechnology). After 5 min, the blot was imaged using a ChemiDoc MP gel imager. These data are shown in **Figure S3**.

**Desthiobiotin Azide Enrichment Procedures.** αS fibrils were treated as described above with irradiation at 254 nm for varying amounts of time. Click chemistry with desthiobiotin azide (Click Chemistry Tools; Scottsdale, AZ, USA) was then conducted as described for fluorophore azides. Samples were then directly applied to 200 μL of streptavidin agarose resin (GoldBio; St. Louis, MO, USA) and allowed to incubate at 4°C overnight. After incubation, the resin was washed 5 times with water, and bound proteins were eluted via treatment with 10 mM biotin (MilliporeSigma, Burlington, MA, USA) for 2 hours at room temperature. Resulting monomeric crosslinked DTBA-labeled αS samples were subjected to a chloroform/methanol/water precipitation (4:3:1), as previously described.[4] The resulting protein pellet was then re-solubilized in 10 μL $H_2O$, 0.1%TFA. The sample was prepared for MALDI-TOF by spotting 1 μL of analyte in 1 μL of sinapic acid matrix (saturated solution in 50:50 $CH_3CN$:$H_2O$, 0.1% TFA) on a ground steel MALDI plate. The sample was then analyzed using linear positive mode with scanning between 5 and 20 kDa at 23% laser power on a Bruker UltraFlex III instrument. These data are shown in **Figure S4**.

***Mass Spectrometry Analysis of α-Synuclein Crosslinking***

**Intact Protein Crosslinking and MALDI-TOF Analysis.** Fibrils were prepared and irradiated in a 1.5 mL tube as described above. After irradiation, 50 µL of 250 mM aqueous SDS solution (250 mM) was added, for a final SDS concentration of 125 mM. Samples were then boiled for 30 min. Resulting monomeric crosslinked αS samples were subjected to a chloroform/methanol/water precipitation (4:3:1), as previously described.[4] The resulting protein pellet was then re-solubilized in 25 µL $H_2O$, 0.1%TFA. The sample was prepared for MALDI-TOF by spotting 1 µL of analyte in 1 µL of sinapic acid matrix (saturated solution in 50:50 $CH_3CN$:$H_2O$, 0.1% TFA) on a ground steel MALDI plate. The sample was then analyzed using linear positive mode with scanning between 5 and 20 kDa at 23% laser power on a Bruker UltraFlex III instrument.

**LC-MS/MS Crosslinking Site Determination.** αS crosslinked peptide samples were prepared as described above for MALDI-TOF analysis. Monomeric αS was re-solubilized in 100 µL ammonium bicarbonate buffer (50 mM $NH_4HCO_3$, pH 7.4) and digested overnight at 37°C via the addition of 1 µg of trypsin (Promega; Madison, WI, USA). Digestion was quenched by adding 1.0 µL of formic acid, and the resulting peptide mixture was cleaned up using custom C18 stage tips.[5] The eluted mixture from the stage tip was then concentrated via Speedvac (Savant), and re-solubilized in 20 µL of $H_2O$ with 0.1% TFA. 2 µL of this sample was then analyzed directly via Orbitrap mass spectrometry on a Thermo QE-HF instrument (see MS/MS Data Acquisition and Analysis section below). These data are shown in **Figure 1** and **Figure S5**.

***Sulfisoxazole and Valdecoxib Crosslinking***

**Cell Culture Procedures.** For routine cultures, MDA-MB-231 or MCF7 cells were grown on 100 mm or 150 mm dishes in sterile-filtered Dulbecco's modified Eagle's medium (DMEM) (Gibco, Thermo Fisher) supplemented with 10% fetal bovine serum (FBS) (Corning), and 1% pen/strep (Mediatech, 10 mcg/ml P – 10 mg/ml S). All cells for stable isotope labeling by amino acids in cell culture (SILAC) experiments were passaged 5 times in SILAC DMEM supplemented with 10% dialyzed FBS and heavy or light lysine ($^{13}C_6{}^{15}N_2$) and arginine ($^{13}C_6{}^{15}N_4$) from a SILAC protein quantitation kit (Thermo Fisher, cat. no. A33972). Cells were then washed with PBS and harvested in 5 ml PBS by scraping. Cells were pelleted by centrifugation at 1000 rpm for 5 min, and the supernatant was removed. The resulting cell pellet was re-suspended in 300-500 μL PBS with 0.1% Triton X-100 (Bio-Rad) and 1x HALT protease inhibitor cocktail without EDTA (Thermo Fisher), then lysed by sonication for two 30 s cycles (cycle 1: 2s on 2s off, amp 50; cycle 2: 2s on 2s off, amp 55) using a QSonica Q700 fitted with a microtip. Following sonication, lysate was centrifuged at 13,200 rpm for 60 min to separate membrane and cytosolic protein fractions. The resulting lysate cytosolic or membrane fraction was then used immediately for photo-crosslinking studies following concentration determination by DC protein assay (Bio-Rad).

**Photo-crosslinking Fluorescence SDS-PAGE Analysis.** Cell lysates were prepared as described above. Samples were then treated with 100 μM probe (**SFX-1** or **VCOX-1**), incubated for 1 h at 37 °C, and irradiated with either 254 nm or 365 nm light (UVP Multiple Ray Lamp, 8 watt) for varying amounts of time. Following irradiation, click chemistry and SDS-PAGE analysis was conducted as described above for αS. These data are shown in **Figure S6** and **S7**.

**Probe Treatments and Enrichment.** MDA-MB-231 or MCF7 lysates were prepared as described above and the concentrations were adjusted to 7.5 mg/mL. Then, 50 μL of the SILAC sample was treated with either 250 μM probe (Heavy "H" condition) or 250 μM probe and 1 mM competitor (Light "L" condition) with a fixed final DMSO concentration of 2%. In each experiment the probe is **SFX-1** or **VCOX-1** and the competitor is the corresponding unmodified compound (**SFX** or **VCOX**). The sample was then incubated at 37 °C for 1 h and irradiated with 254 nm light for 10 min (Analytik Jena UVP Multiple Ray Lamp, 8 watt). Following irradiation, samples were conjugated to a UV-cleavable biotin azide probe (Click Chemistry Tools) via copper-catalyzed click chemistry as described above for αS. Heavy and light lysates were then diluted with 400 μL PBS + 0.05% tween20, mixed, and applied directly to 200 μL of streptavidin-agarose resin (GoldBio). Samples were allowed to rotate at 4 °C overnight. The samples were then centrifuged at 2000 rpm for 3 min, and the flowthrough was saved. The resulting slurry was washed 5 times with PBS + 0.05% tween20, and 3 times with LC-MS grade water (Pierce Biotechnology). The washed resin was then resuspended in 500 μL of LC-MS grade water and irradiated at 365 nm for 30-45 min with shaking to fully cleave the enriched proteins from the resin. Note that in our studies of isoxazole irradiation under different wavelengths of light, reactions were negligible with 365 nm irradiation. The eluate was then harvested via centrifugation, and the resin was washed again with 500 μL of water and collected. The eluates were then combined and concentrated via speed-vacuum (Savant).

*MS/MS Data Acquisition and Analysis*

**General Procedure for LC-MS/MS Experiments.** Dried samples were resuspended in 50 µL of triethylammonium bicarbonate (TEAB) resuspension buffer (2.5% SDS and 50 mM TEAB final concentrations) and reduced with final 10 mM DTT (US Biological; Salem, MA, USA) for 30 min at 30 °C, followed by alkylation with final 50 mM iodoacetamide (Sigma Aldrich) for 30 min at 30 °C. The proteins were captured in an S-TrapTM mini column (C02-mini, Protifi; Farmingdale, NY, USA) to remove contaminants, salts, and detergents and concentrate the proteins in the column for efficient digestion then digested with trypsin (Thermo Fisher Scientific) in 1:10 (w/w) enzyme/protein ratio for 1 h at 47 °C. Peptides eluted from this column, with 50 mM TEAB, 0.2% formic acid, and 60% acetonitrile in order, were vacuum-dried and resuspended with 0.1% (v/v) TFA in LC-MS grade water for mass spectrometry analysis.

Digested samples were analyzed by a Q-Exactive HF mass spectrometer (Thermo Fisher Scientific) coupled to a Dionex Ultimate 3000 UHPLC system (Thermo Fischer Scientific) equipped with an in-house made 15 cm long fused silica capillary column (75 µm ID), packed with reversed phase Repro-Sil Pure C18-AQ 2.4 µm resin (Dr. Maisch GmbH, Ammerbuch, Germany). Elution was performed by the following method: a linear gradient from 4 to 38% buffer B (90 min), followed by 95% buffer B (5 min), and re-equilibration from 95 to 4% buffer B (5 min) with a flow rate of 300 nL/min (buffer A: 0.1% formic acid in water; buffer B: 80% acetonitrile with 0.1% formic acid). Data were acquired in data-dependent MS/MS mode. Full scan MS settings were as follows: mass range 200–1600 m/z, resolution 120,000; MS1 AGC target 3E6; MS1 Maximum IT 100. MS/MS settings were: resolution 30,000; AGC target 5E5; MS2 Maximum IT 100 ms; fragmentation was enforced by higher-energy collisional dissociation with stepped collision energy of 25, 27, 30; loop count top 20; isolation window 1.4; MS2 Minimum AGC target 800; charge exclusion: unassigned, 1, 8 and >8; peptide match preferred; exclude isotope on; dynamic exclusion 45 s.

**MS/MS Data Analysis. AS3** αS crosslinking samples were searched using Proteome Discoverer (Thermo Fisher) with Sequest HT against the *E. coli* proteome with αS sequence information added in. Precursor mass tolerance was set to 10 ppm and fragment mass tolerance was set to 0.02 Da. **AS3** crosslinker was specified as a dynamic modification of +354.158 Da on all amino acids.

Search parameters were set to full trypsin digestion, with maxima of four missed cleavages and three modifications per peptide. Acetylation, Met-loss, and Met-loss+acetylation were also specified as dynamic modifications. A strict false discovery rate (FDR) of 0.01 was used for the identification of high-confidence targets and a relaxed FDR of 0.05 was used for differentiation of medium and low-confidence targets. Following database searching, peptides identified as belonging to αS were filtered for containing the **AS3** modification as well as not being present in the DMSO control sample. The peptides were then sorted by descending raw abundance values, and spectra were each manually inspected to confirm crosslinking.

Briefly, for **SFX/VCOX** comparison data, proteins were filtered for those containing greater than or equal to 20 peptide spectral matches and were analyzed for those enriched (>1.5 fold change) in **SFX** pulldowns and un-enriched in **VCOX** pulldowns (<1.5 fold change). This qualitative analysis provides a general measure of the specificity of **SFX** or **VCOX** for protein subsets. These data are shown in **Figures S8** and **S9**. Filtered and unfiltered spreadsheets of all **SFX/VCOX** protein targets can be found in the associated file **SFX_VCOX_Comparisons.xlsx**.

**SFX-1**/**SFX-1** SILAC samples were searched using MaxQuant (Version 1.6.17.0) against the reviewed human proteome (accessed 11-15-2021). A multiplicity of 2 was used with Arg10 ("Heavy" Arginine, $^{13}C_6{}^{15}N_4$) and Lys8 ("Heavy" Lysine, $^{13}C_6{}^{15}N_2$) specified for quantification, with a maximum labeling setting of 3. Met oxidation, N-terminal acetylation, and carbamido-methylation were specified as dynamic modifications. All other settings were default using the "orbitrap" instrument type. The first search peptide tolerance was set to 20 ppm and the main search peptide tolerance was set to 4.5 ppm. Under "*miscellaneous*", re-quantify was checked to bolster quantification of peptides identified containing only one labeling pair. Under protein quantification in global parameters, the label minimum ratio count for quantification was set to 1.

Following identification and quantification, several python scripts were used to analyze data. Briefly, minimalist filtering was conducted to remove contaminants, reverse hits, and proteins with quantification in less than 70% of replicates. A k-Nearest Neighbor (kNN) imputation approach was then adopted using the sklearn module in Python, with optimization of k against a dummy dataframe constructed by arbitrarily removing values from proteins with quantification across all replicates to match the percentage of missing data in the real dataset. The root mean square error (RMSE) of the k-imputed dummy dataset can then be cross-validated against the real dataset from which it was derived to optimize k. After this, a one-sample t-test against a theoretical population mean of 1 (no enrichment in SILAC experiment) was conducted to gauge confidence in enrichment of the given proteins, and the Benjamini-Hochberg correction was implemented with an FDR of 0.01. More stringent filtering was then conducted to select for proteins with a computed H/L ratio of greater than or equal to 1.5, and an andromeda score of greater than 15 (just above medium confidence). This filtered protein list is available as a supplement to this manuscript. The highly enriched protein dataset was then annotated according to families and domains present in the Uniprot database, and the annotated domains were amalgamated to reduce redundancies. The domain list was then one-hot encoded and filtered for the top-10 most abundant domains by number of proteins associated. The resulting protein list was then clustered using k-means with optimization of k using the silhouette coefficient. The resulting clusters were then analyzed for the domains present. These data are shown in **Figure 2**. Filtered and unfiltered spreadsheets along with quantification and clustering of all **SFX** protein targets can be found in the associated file **SFX_Proteomics_Data.xlsx**.

**Calmodulin Crosslinking Verification.** CaM was crosslinked and analyzed by SDS-PAGE in-gel fluorescence similarly to αS. CaM (100 µM) prepared as described above was treated with 100 µM **SFX** probe molecule from a DMSO stock (50 µL total volume, 1% DMSO) in a 1.5 mL tube. The sample was incubated at 37 °C for 1 h. Following incubation, the lid was opened, and the sample was irradiated with shaking (500-750 rpm) for 10 min, with the lamp (UVP Multiple Ray Lamp, 254 nm, 8 watt) positioned approximately 1-2 cm from the top of the tube. Following irradiation, CaM was directly subjected to copper-catalyzed click chemistry. Briefly, 16.5 µL of PBS was added to each sample to pre-adjust for the final reaction volume. 1.0 µL of TAMRA-azide (Lumiprobe) was then added to each sample. Click-mix was made by pre-mixing 1.875 µL $CuSO_4$ (25 mM) with 1.875 µL THPTA (50 mM) per reaction/sample (i.e. 10 samples = 18.75 µL of each). 3.75 µL of this Click-mix was then charged to each sample ($CuSO_4$ final concentration: 500 µM, THPTA final concentration: 1.25 mM). Finally, 3.75 µL of sodium ascorbate (40 mM) was added (final concentration: 2 mM). Reactions were allowed to react with gentle agitation at room temperature for 1.5-2 h. Reactions were then quenched via the addition 25 µL 4xLDS loading buffer (Thermo Fisher) with dithiothreitol (DTT, 200 mM) and were heated at 75 °C for 10 min. 20 µL of the resulting solution was then separated via SDS-PAGE and imaged for fluorescence on a Chemidoc gel imager (Bio-Rad).

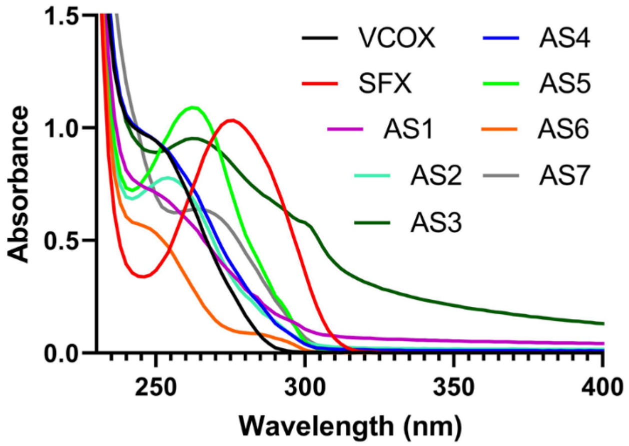


**Figure S1.** UV-Vis spectra of all photo-crosslinking probes at 100 μM in 50/50 MeOH:Water obtained on a Thermo Fisher Scientific GENESYS 150 UV-Vis spectrophotometer.

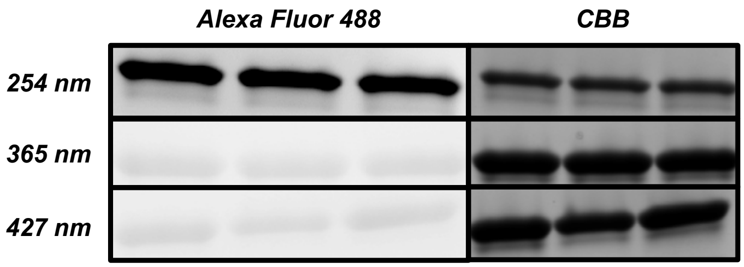


**Figure S2.** Photo-crosslinking of αS with **AS3** under irradiation by varying wavelengths of light.

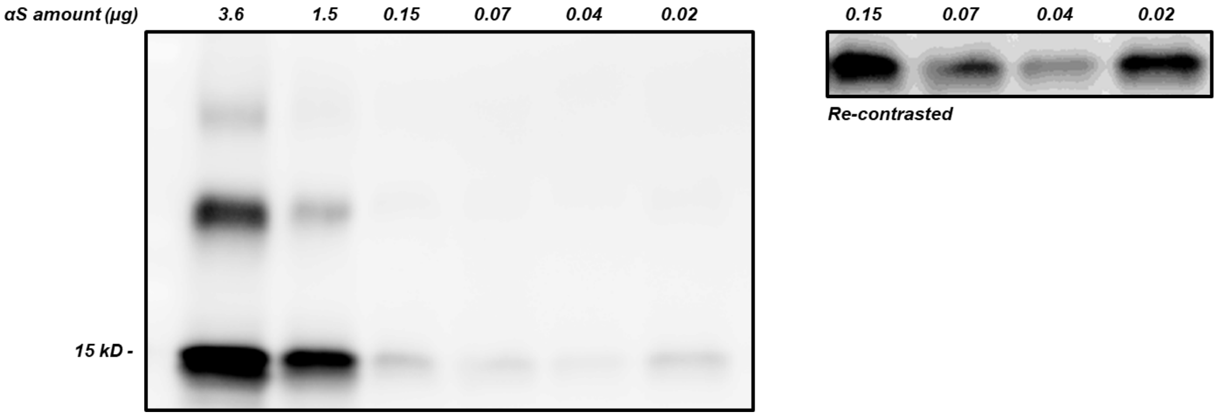


**Figure S3.** Assessment of the sensitivity of probe **AS3** in identifying various amounts of αS via enhanced chemiluminescence (ECL) blotting.

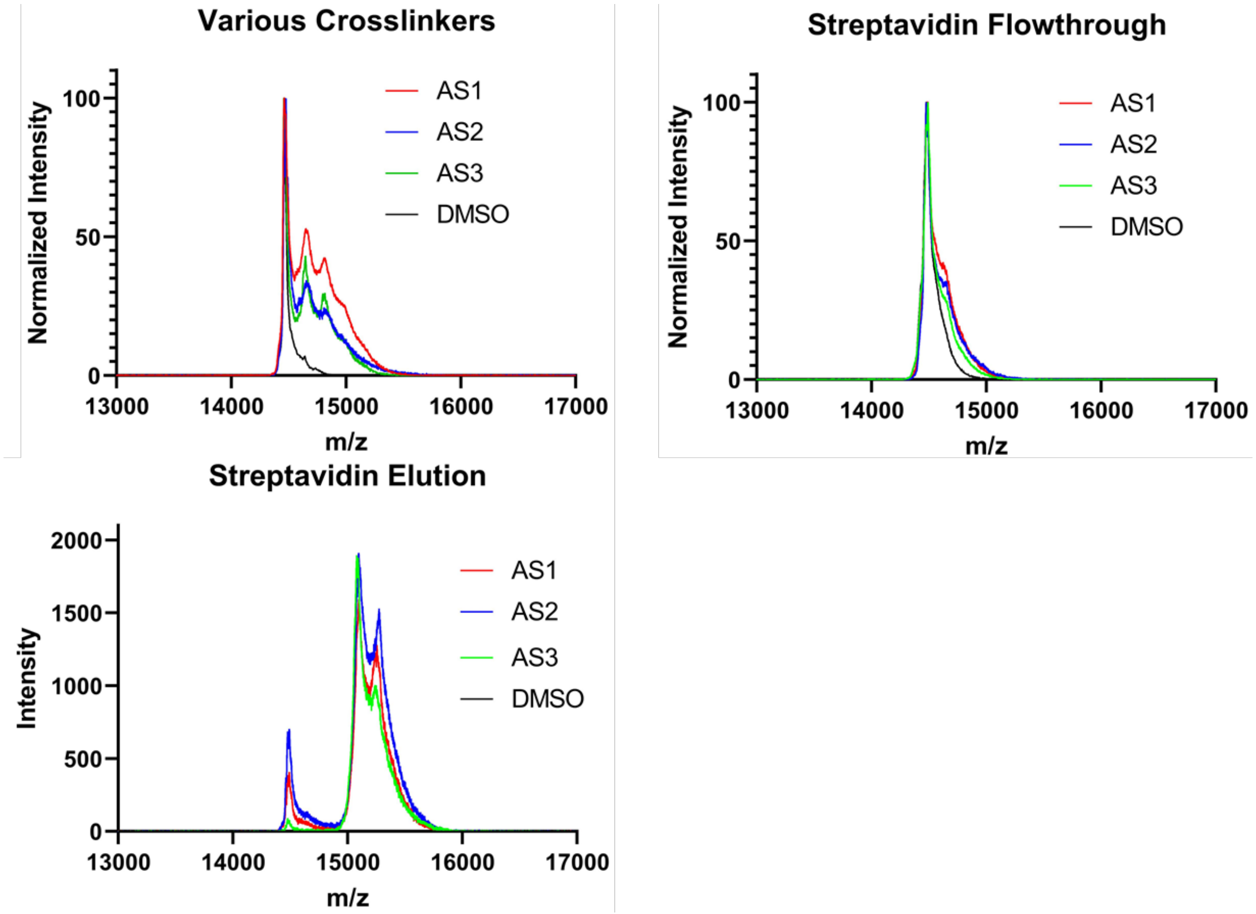


**Figure S4.** Validation of **A**. crosslinking **B**. depletion by streptavidin and **C**. elution from streptavidin of various photo-crosslinkers via click chemistry with desthiobiotin azide and subsequent enrichment.

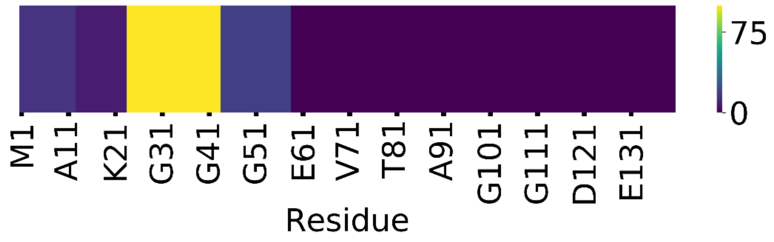


| ***Annotated Sequence*** | ***Positions in Master Proteins*** | ***Normalized Abundance*** |
|---|---|---|
| [K].QGVAEAAGKTKEGVLYVGSK.[T] | [24-43] | 100.00 |
| [K].EGVLYVGSKTKEGVVHGVATVAEK.[T] | [35-58] | 18.45 |
| [-].MDVFMKGLSKAK.[E] | [1-12] | 15.16 |
| [K].AKEGVVAAAEKTKQGVAEAAGK.[T] | [11-32] | 7.59 |
| [K].TKEGVLYVGSK.[T] | [33-43] | 4.20 |

**Figure S5. AS3** photo-crosslink enabled binding site mapping in αS.

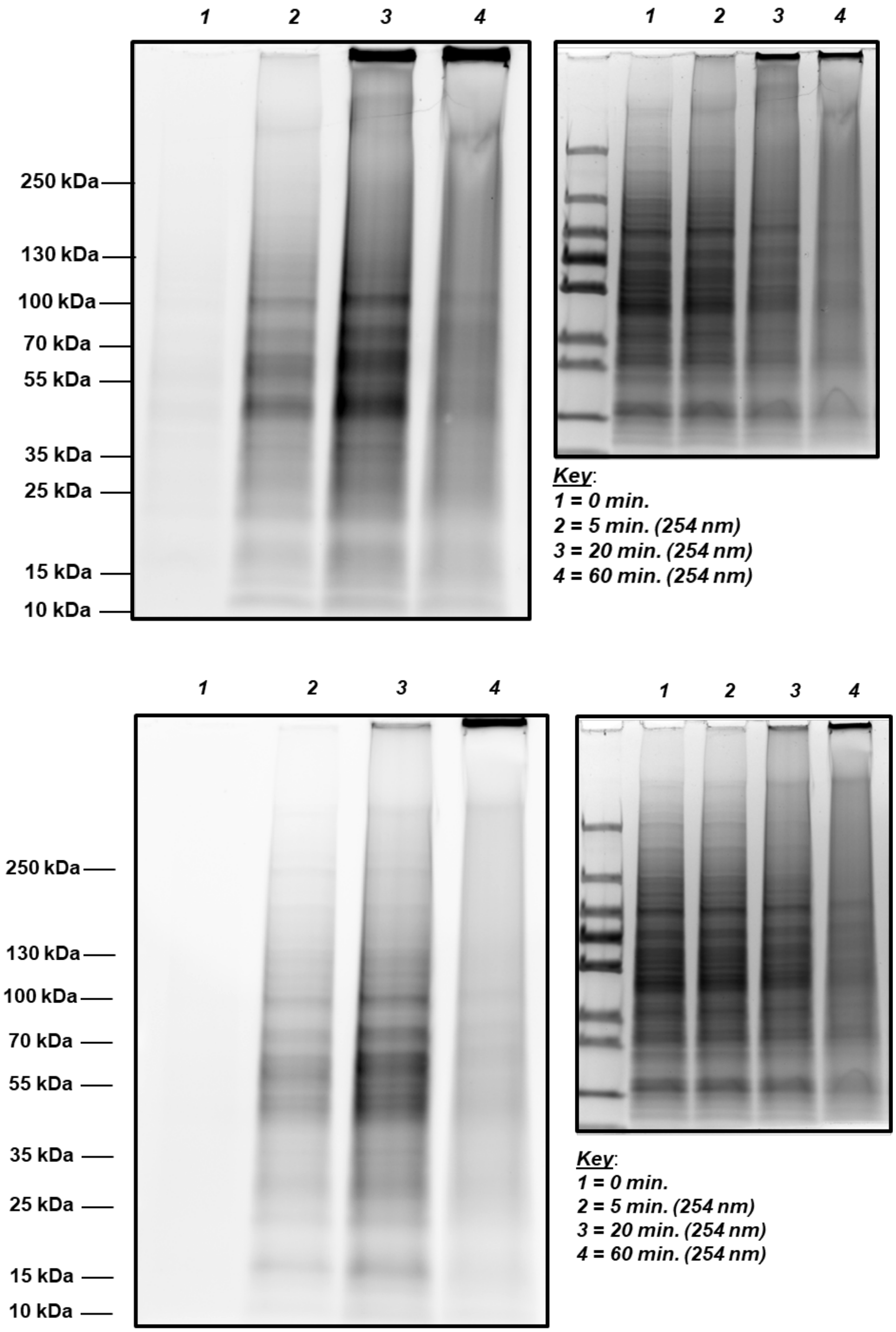


**Figure S6.** Time-dependent light response of top - sulfisoxazole probe (**SFX-1**) in the presence of HEK cell lysate (2 mg/mL) and bottom – **VCOX-1**. Substantial aggregation can be observed at the top of the gel following irradiation times longer than 20 min.

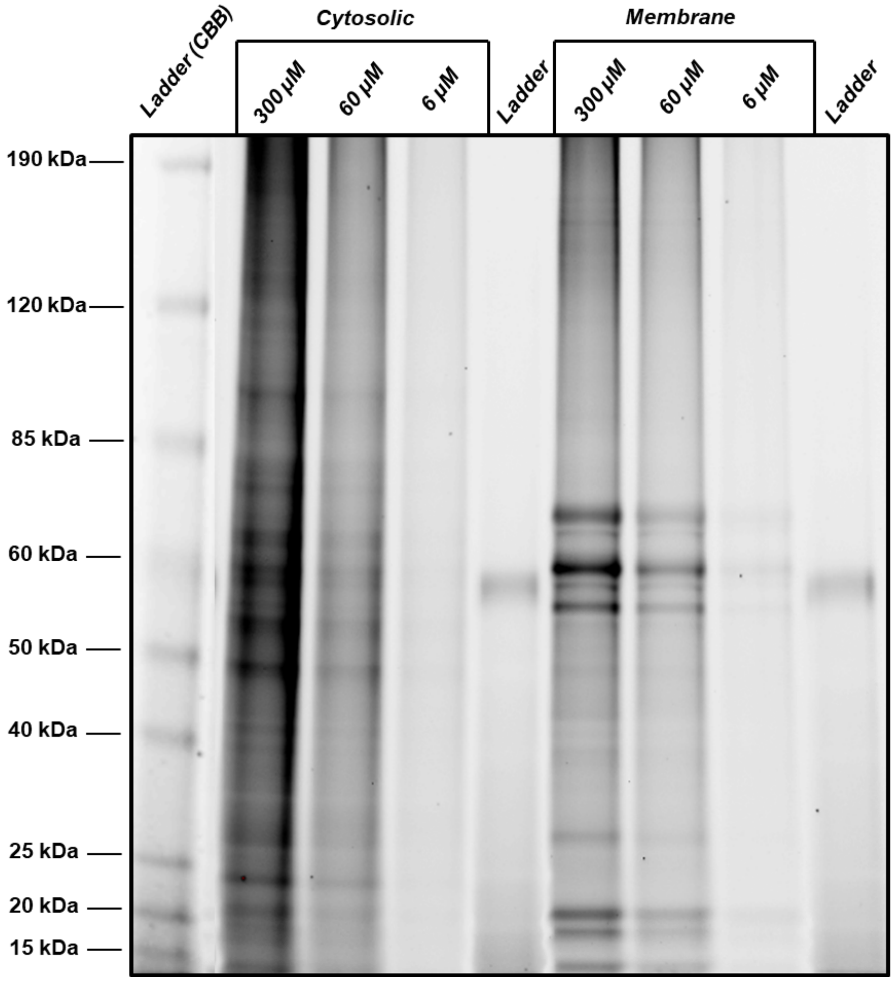


**Figure S7.** Photo-crosslinking and TAMRA-SDS-PAGE of **SFX-1** in MCF7 cells demonstrating covalent labeling of multiple proteins with a distinct dose response.

| *SFX* | *VCOX* | *Both (SFX-ranked)* | *Both (VCOX-ranked)* |
|---|---|---|---|
| CALM | ALDOC | SLC25A4 | SLC25A4 |
| RDX | VIM | MYH9 | TUBB8 |
| MSN | KRT18 | RPL4 | LMNA |
| GNS | KRT8 | PPP1CC | PLEC |
| GSS | ACTN1 | TUBB4A | LMNB1 |
| TPI1 | ACTN4 | RPS11 | TUBA1C |
| ENO2 | KRT19 | RPS8 | EIF3CL |
| TBCA | KRT7 | TUBB6 | LRPPRC |
| ANXA5 | SPTBN1 | RPS4X | PABPC4 |
| HSD17B4 | IMPDH2 | RPS3A | TUBB4A |
| BTF3 | LMNB2 | TUBB8 | EIF3L |
| S100A6 | NUMA1 | EIF4A2 | MCM2 |
| STMN1 | TPM1 | RPS16 | MYH9 |
| MAT2A | SEPT7 | DYNC1H1 | DYNC1H1 |
| HIST2H2BF | TPR | TUBA1A | CCT2 |
| PARK7 | KTN1 | DDX5 | CCT5 |
| HIST1H4A | TGM2 | ARF4 | EIF3A |
| RBM3 | PAICS | TUBA1C | PPP1CC |
| DPP3 | CCT8 | TUBA1B | PSMD12 |
| NCL | RTN4 | XPO1 | CCT4 |

**Figure S8.** Table of top 20 protein hit gene names in each probe subset ranked in order of H:L ratio.

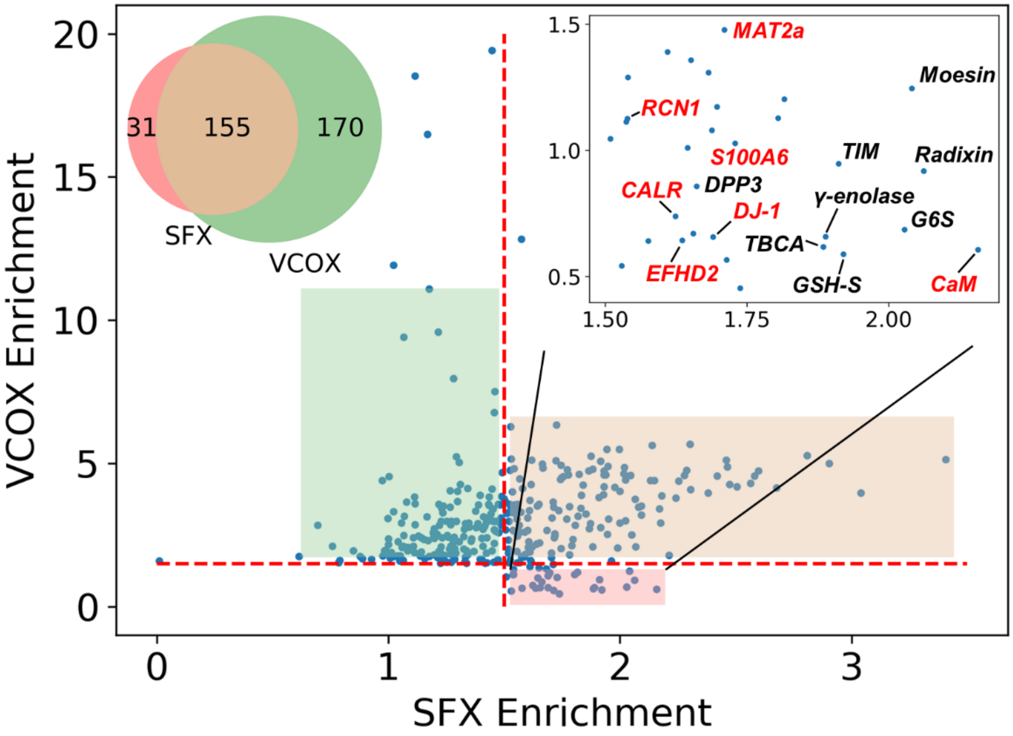


**Figure S9.** Two-dimensional enrichment profiles of proteins identified in **SFX/VCOX** cross comparison SILAC studies, plotted as the heavy/light ratio obtained in **VCOX** samples vs. the H:L ratio obtained in **SFX** samples. Areas highlighted in green, red, or brown refer to proteins enriched in **VCOX**, **SFX**, or both respectively.
